# A high-resolution microscopy system for biological studies of cold-adapted species under physiological conditions

**DOI:** 10.1101/2024.09.17.613460

**Authors:** Anne-Pia M. Marty, Edward N. Ward, Jacob R. Lamb, Francesca W. van Tartwijk, Lloyd S. Peck, Melody S. Clark, Clemens F. Kaminski

**Affiliations:** University of Cambridge, Department of Chemical Engineering and Biotechnology, Phillipa Fawcett Drive, Cambridge CB3 0AS United Kingdom; British Antarctic Survey, High Cross, Madingley Road, Cambridge CB3 0ET United Kingdom

**Author notes:** Corresponding author: Clemens F. Kaminski. MRC Laboratory of Molecular Biology, Francis Crick Avenue, Cambridge Biomedical Campus, Cambridge CB2 0QH, UK.

**Keywords:** Optical microscopy, Super-resolution, Temperature, cold-adaptation, extremophiles

## Abstract

The fundamental processes governing life are sensitively dependent on temperature. Whilst much is known about the constraints on how proteins operate at 37°C, little knowledge exists about how biological function is maintained sub-zero temperature conditions, where proteins are less stable and oxidative damage is high. However, almost 90% of habitable environments on Earth are permanently below 5°C (i.e. the deep sea and polar regions). This means that we do not understand how a large and diverse proportion of the global biome functions. To address this question at the cellular level, tools are required for imaging biological systems at high resolution under physiological conditions. This poses severe technical challenges that cannot be addressed with traditional optical microscopy techniques. High-resolution imaging objectives require short working distances and the use of immersion media, which lead to rapid heat transfer from the microscope to the sample. This affects the viability of live specimens and the interpretability of the results when the sample function optimally at low temperatures. Condensation and temperature-induced shrinking of components pose further challenges, reducing image resolution and contrast. Here, we address these issues and provide a method for high-fidelity imaging of live biological samples at temperatures of around, or below, 0°C. Our method is compatible with different microscopy modalities, including super-resolution imaging. It relies on hardware additions to traditional microscopy systems that can be straightforwardly implemented, namely, a cooling collar, 10% ethanol as an immersion medium, and nitrogen flow to mitigate condensation. We demonstrate the method in live cell cultures derived from Antarctic fish species and highlight the need to maintain physiological conditions for these fragile biological samples. Future applications are diverse and include evolutionary biology and the study of cold-adapted organisms, as well as cellular biophysics and several applications in biotechnology.

## 1. Introduction

The fundamental processes of life are governed by elementary chemical and physical mechanisms, such as diffusion, transport, and chemical reactions, all of which depend sensitively on temperature. At different scales, from molecular to cellular levels, temperature also impacts density, viscosity, gas solubility, etc. Since these physical properties are interdependent, and all are governed by temperature, most life forms have evolved to operate over their respective optimal temperature ranges, outside of which they cannot survive [1].

While biological processes at normothermic conditions (e.g., 37°C in human physiology) are well-characterised, the quality of model predictions are often less good for organisms with lower normothermic temperatures. This decline in quality is particularly evident at the Arrhenius Break Temperature (ABT), which typically ranges between 2°C and 5°C, depending on the specific biological process [2], [3]: below the ABT, the relationship between the logarithm of biochemical reaction rates and the inverse of temperature deviates from linearity, indicating a fundamental change in the rate-limiting steps driving these reactions. This deviation is evident also from the disproportionately increasing timescales below the ABT for life-sustaining processes like protein folding, cell division, and animal development.[2]

Research into the biophysical adaptations of life at temperatures around 0°C is not merely of fundamental academic interest but is of ecological relevance for understudied environmental conditions. Notably, over 88% of the oceanic volume lies at depths exceeding a thousand metres, beneath which the temperatures hover around 4°C [4]. Contrary to cold habitats on land, where direct exposure to the sun can make temperatures rise by tens of degrees, the thermal inertia of water makes marine temperature changes minimal. The Southern Ocean’s surface temperatures range between -2°C and 2°C all year round [3]. The vast and diverse biome of cold oceans is home to numerous species whose survival strategies and basic life mechanisms remain poorly understood, due to the inadequacy of existing models describing life at such low temperatures. Much of this biome is at risk from global warming and rising water temperatures, and efforts to improve understanding of cold adaptation are therefore both timely and important.

Research into cold-adapted biological systems has thus far been limited by the fact that traditional biotechnological tools, including various forms of microscopy, are not designed to function well at low temperature. In the case of microscopy, image quality suffers, and it is difficult to maintain cold-adapted samples at their physiological temperature. In this work, we address these challenges, developing methodologies and tools capable of accurately probing biological phenomena at the subcellular level at near-freezing conditions.

Achieving cooling at the focal plane of a microscope presents several issues, such as ice formation, condensation, and sample heating (Figure 1). One could house an entire microscope system in a cold room, but this restricts the use of expensive equipment to one type of application only. It also makes changing temperature conditions difficult and slow, as thermal inertia means hours of settling time are required. Condensation problems can also occur during warming up or cooling down of microscope components as they transition through atmospheric dew points. This is especially critical during maintenance or servicing of cold laboratories, when the temperature is brought to ambient levels. Finally, using samples in aqueous solutions or simply the presence of experimenters near the apparatus introduce humidity which will freeze over any exposed surface.

**Figure 1.**
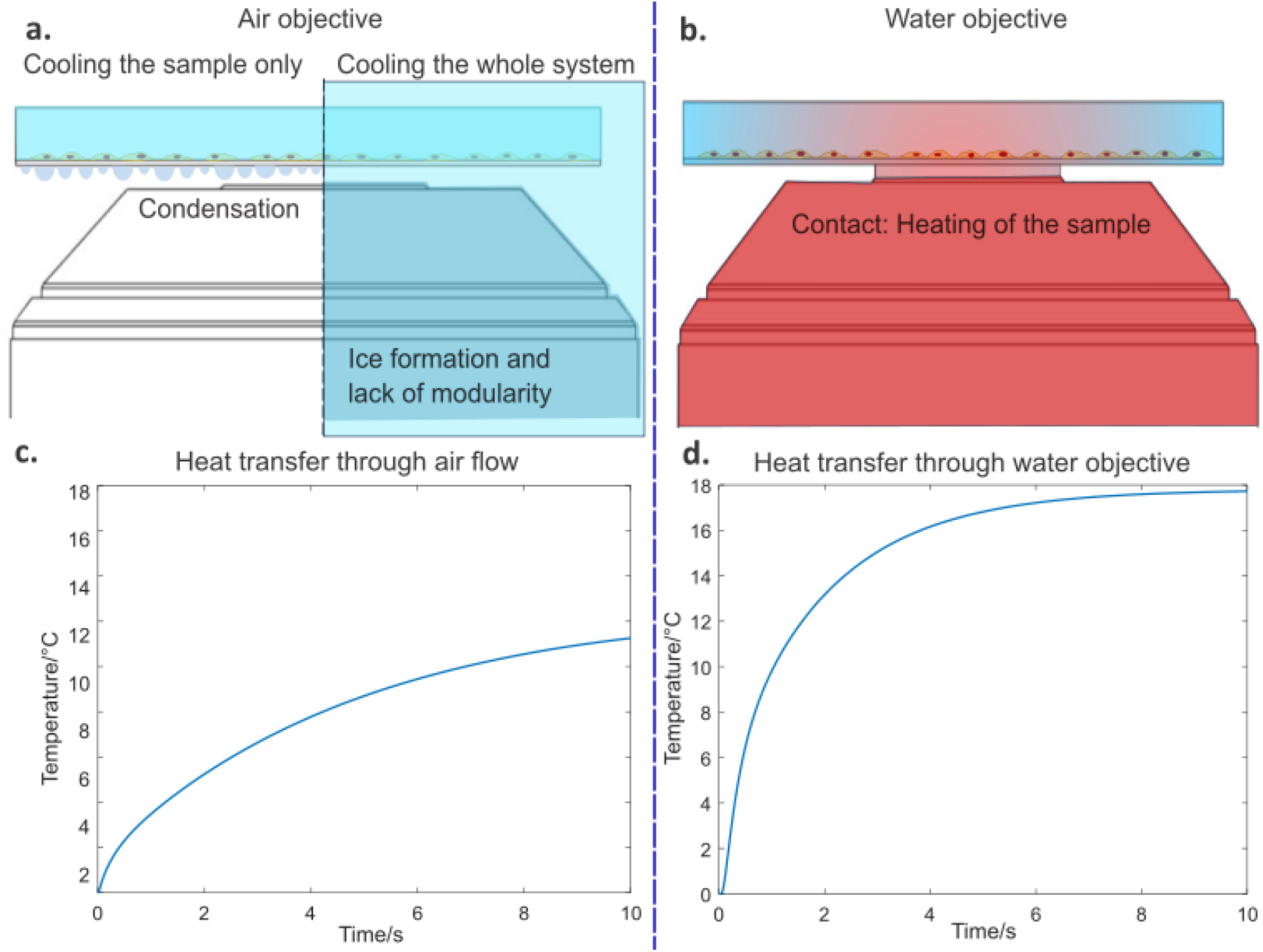
Heat transfer poses challenges during imaging of samples at temperatures significantly below ambient. a) Left: Using an air objective while keeping the sample cold but the microscope at ambient temperature leads to condensation on optical surfaces. Right: Condensation is prevented by operating the microscope in a cooled environment; however, the risk of ice formation severely limits operational freedom. b) Using a liquid-immersion objective maintained at ambient temperatures heats the sample through thermal contact. c) and d): Simulations show that due to the very short working distance of high-resolution objective lenses, heat transfer through either convection – with an air immersion lens – or conduction – through a liquid immersion lens – rapidly heats the sample at the focal plane of the imaging lens. In both simulations, the sample chamber is cooled externally, but the imaging lens remains at ambient temperatures. More details on simulation parameters are given in S1.

Fully enclosed sample chambers [4], [5], [6] were used to prevent condensation in previous work, but this has critical disadvantages, relying on low resolution widefield microscopy [7], or requiring cooling without simultaneous high resolution imaging [8], [9]. These chambers introduce additional glass surfaces between the sample and the imaging lens, increasing working distance and thus requiring the use of low numerical aperture (NA) lenses. Furthermore, optical mismatches introduce spherical aberrations and lower the image quality.

Using a cooling stage rather than an enclosed chamber, on the other hand, introduces an air gap between the imaging lens and the cooled sample and this inevitably generates condensation (Figure 1a). It is not easily possible to mitigate this problem. For example, attempting to prevent condensation by blowing a stream of dry gas into the space between the sample and the objective results in the sample quickly reaching the air temperature via convective heating. A simulation of this effect is shown in Figure 1c. Further details on the simulation method are provided in Figure S1 and Table S1. The use of an immersion medium between the sample and imaging lens circumvents the condensation problem associated with cooling stages, but massively increases heat conduction from the objective to the sample. This leads to the sample almost immediately reaching the temperature of the lens as shown in the simulation presented in Figure 1d. Therefore, existing solutions such as cooling stages can be useful to keep the sample cold but cannot simultaneously achieve conditions for high-quality imaging *and* the maintenance of low temperatures within the field of view.

To address this, we propose a design for a cooling chamber (Figure S2) and propose modifications to standard optical microscopes that permit the imaging of samples whilst maintaining them at their physiological temperature around 0°C. The methods presented here prevent issues of condensation and heat conduction and maintain image resolution whilst keeping fragile live samples viable.

## 2. Materials and Methods

### 2.1 Thermal simulations

Heat transfer simulations were performed using the MATLAB PDE toolbox, generating a model of the objective to sample contact with the dimensions and physical values as stated in Figure S1 and Table S1, respectively. The model was run for 30,000 steps, with 1000 steps/second.

Simulations for the thermal performance of the collar are shown in Figure S3. Calculations were performed using the Fusion360 thermal simulation workspace. All .obj files, code scripts and simulation parameters are available on github.com/piamrt/coolmicroscope. Thermal loads were set to 20°C air temperature, and -15°C collar temperature.

### 2.2 Refractive index measurements

Refractive indices were measured using an Abbe ‘60’ Refractometer from Bellingham & Stanley Limited. The refractometer was coupled to a Cole Palmer Polystat 12104-05 water chiller and pump system, circulating a 70:30 water-glycerol mix. The light source was a 4-wavelength LED source (LED4D245 Thorlabs) with a DC4104 LED driver.

### 2.3 Hardware design

The collar was manufactured from custom specifications by the company Instec. The collar holder and the air inlet were designed using the Fusion360 software. The resulting Obj output files were sliced using the Ultimaker Cura software and printed in PLA (Ultimaker) on Ultimaker printers. All design files are accessible on github.com/piamrt/coolmicroscope.

The sample chamber shown in Figure S2 was also designed with Fusion 360 and machining steps were defined in the manufacturing mode of the software. The chamber was machined from aluminium with a Siemens XYZ Machinetools XYZ 500LR CNC mill. The chamber was designed to accommodate Lab Tech 8- and 4-well chambers, as well as ibidi µ-slides and 35 mm round culture dishes.

### 2.4 Temperature and power measurements

8-well 1.7 mm glass-bottom cell-culture well-plates (Lab-Teck) were filled with 500 µL water to simulate culture medium. In-well temperatures were recorded using a TC-08 thermocouple data logger (Piotech). The probes were type K exposed junction thermocouples with a tip diameter of 1.5 mm. Small holes were pierced into the lid of the culture chambers. Probes were attached to the wells to be in contact with the bottom of the well and the microscopes were focused so that the tip of the probe was in focus. Data were acquired in continuous data acquisition mode at a sampling rate of one datapoint per second. The power of the laser was measured with a Thorlabs PM100D Power Meter and an S130C probe (Thorlabs). Results presented correspond to the average of 5 repeat measurements.

### 2.5 Image processing

Monolayers of 0.1 µm diameter TetraSpeck beads (ThermoFisher) were produced using a 10^−4^ dilution of the stock solution. Fluorescence image stacks of the beads were recorded over the field of view, with a step size of 0.02 µm along the optical axis. The code for generating image stacks, segmenting the beads, and calculation of their full width at half maximum (FWHM) is available on github.com/piamrt/coolmicroscope.

### 2.6 Super-resolution imaging

Structured Illumination Microscopy (SIM) was the system used for super-resolution imaging. The microscope was built according to the instructions in [10]. SIM is a method relying on the generation of Moiré Patterns from the interference between the sample spatial frequencies and the patterning generated in the excitation light. Super-resolution images are reconstructed from analysis of the patterns. This method typically doubles the resolution of diffraction-limited microscopy. Our SIM system uses a IX71 microscope stage (Olympus) and three laser sources for excitation: 488 nm (iBEAM-SMART-488, Toptica), 561 nm (OBIS 561, Coherent), and 640 nm (MLD 640, Cobolt). Patterning was done with a ferroelectric binary Spatial Light Modulator (SLM) (SXGA-3DM, Forth Dimension Displays) and polarisation was controlled with a Pockels cell (M350-80-01, Conoptics). The camera used was an sCMOS camera (C11440, Hamamatsu), the raw image acquisition and recording was managed by the HCImage software (Hamamatsu) and the hardware was synchronised using a custom LabView program (available upon request).

### 2.7 Fixed samples

Fixed cells were used to compare the quality of imaging across a temperature range, on a biological sample that would not dynamically change with temperature. Samples of mammalian (Vero) cells were fixed using 4% Paraformaldehyde (PFA) and 0.1% glutaraldehyde in PBS for 10 minutes at room temperature. Cells were permeabilised using Triton x-100 0.5% in PBS also for 10 minutes at room temperature. The subsequent blocking step used 10% goat serum in PBS for 30 min at room temperature. The Primary antibody was a Mouse AB13120 β-tubulin-targeting antibody and the secondary antibody was Rabbit AB6046 (Abcam). Both were diluted 1:400 in PBS with 2% Bovine Serum Albumin (ThermoFisher) and 0.005% Triton, and each was incubated 1h at room temperature, the last step being performed in the dark.

### 2.8 Antarctic fish cell culture and imaging

Primary cell cultures were obtained from dissected tissues of *Harpagifer antarcticus* (unpublished methods). All *H. antarcticus* used in the experimental work were collected at Rothera Research Station, Adelaide Island, Antarctic Peninsula (67°34′07″ S, 68°07′ 30″ W) by SCUBA divers during the austral summer. The fish were returned to the UK and maintained in a recirculating aquarium at temperatures close to 0 °C until required. Fish were collected under a permit (BAS-S7-2022/01) granted under Section 7 of the Antarctic Act 1994. Before dissections were performed for cell cultures, all fish were killed according to Home Office UK schedule 1 requirements. Cell monolayers were stained with MitoTracker green according to the manufacturer recommendations. Samples were imaged at 2°C, brought to 20°C for 1h and then imaged again at 2°C using a structured illumination microscope (SIM) [11].

TIF files were reconstructed using the Fiji LAG-SIM package, available from the Fiji Update sites. LAG SIM runs on the fairSIM [12] package and provides an interface allowing iteration through different parameters and processing of batches of data.

The contrast of the reconstructed images was enhanced manually, using the Fiji image processing tools ‘Enhance Local Contrast’ and ‘adjust Brightness/Contrast’ and the images were exported in JPEG format to be analysed in CellProfiler[13]. The following CellProfiler pipeline was used to analyse the mitochondrial phenotype: ColorToGray > RescaleIntensity > MedianFilter >IdentifyPrimaryObjects > MeasureObjectSizeShape > ExportToSpreadsheet > OverlayObjects > SaveImages.

The resulting CSV files were then imported into MATLAB for plotting and to perform Student’s t-tests.

## 3. Results and Discussion

Our design was informed by model predictions of a high rate of heat transfer between the objective lens and the sample through the immersion medium (water or 10% ethanol; Figure 1d). We decided to use this phenomenon for maintaining the correct temperature of the sample around the focal volume. A cooling element was attached close to the tip of the objective (Figure 2a). Its purpose was to cool the objective to a temperature of around -15° C and for the objective to act as a heat sink with high thermal mass. Subsequent heat transfer from the sample to the objective reduced the sample temperature to the desired range of around 0°C (Figure 2b). Thermal simulations show that a cooling the collar must theoretically be maintained at -20°C to allow the temperature of the sample to reach -6°C at the point of focus (full details are given in Figure S3 and corresponding text). Variations in objective shapes and size mean that the use of different objective models requires new calibration curves to be established. Figure 2b shows data for a 60x water objective (Olympus UPLSAPO60XW, used for all experiments presented here). TC-08 thermocouple probes (PicoTech) were used in an 8-well glass-bottom (0.17 mm tick) dish to record the temperature near the focal point of the objective, verifying that sub-0°C temperatures are reachable in the sample volume.

**Figure 2.**
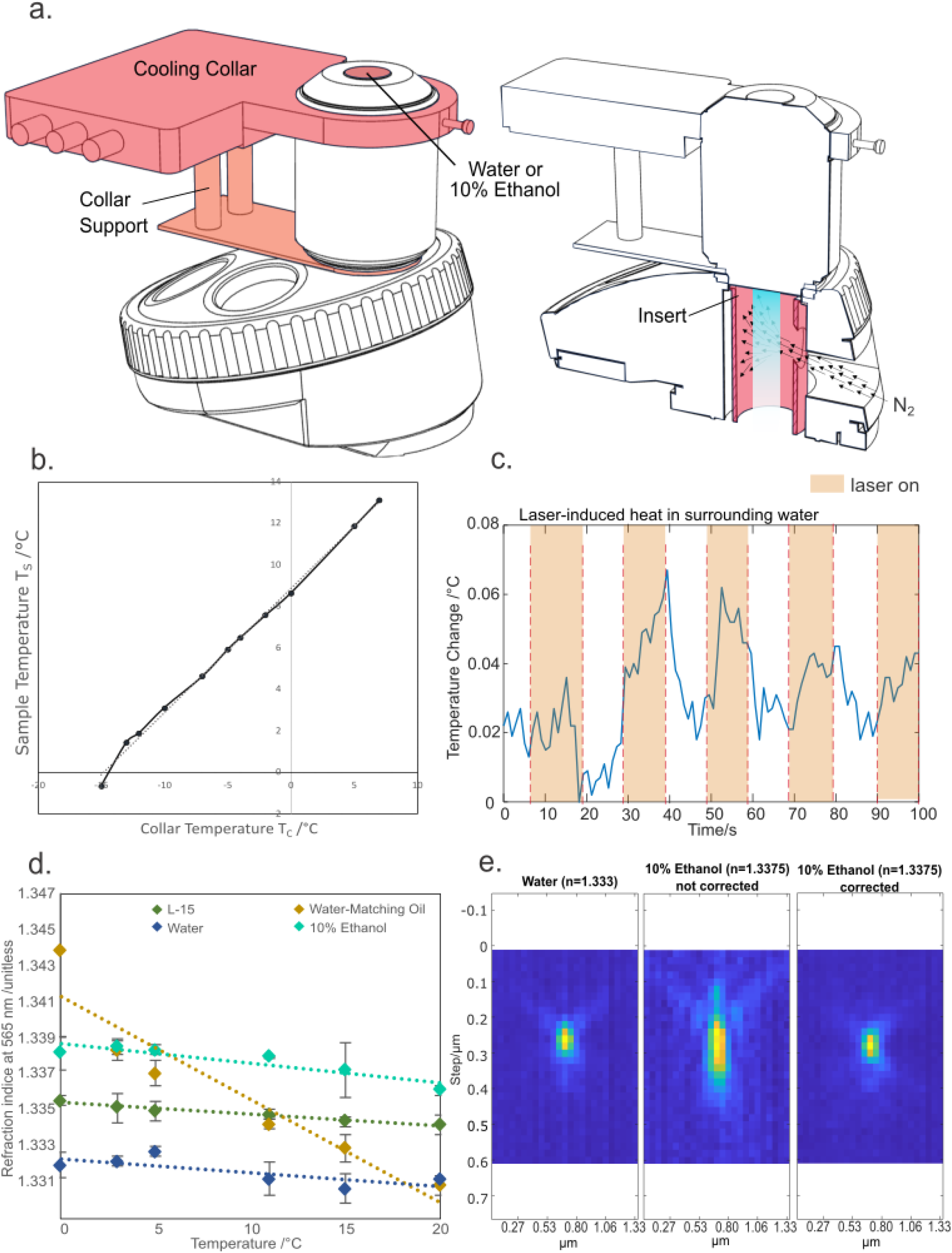
The use of a cooled water-immersion lens allows for imaging at sub-zero temperatures with no loss in image resolution. a) Left: A supported cooling collar attached to the top of the objective keeps the image plane at the desired temperature while the microscope remains at room temperature. Right: To prevent condensation of water at the rear of the objective, the air behind the objective lens is purged with dry gas. b) Temperature reached at the focal volume plotted as a function of the collar temperature. Straight line fit: T_S_=0.59T_C_ + 8.87. c) Heat generated by the laser excitation resulted in a temperature change of <0.1°C. d) Temperature dependent variation in refractive index for objective immersion liquids. Optimal 3D imaging of samples requires an objective immersion liquid with a refractive index equal to that of the sample (L-15 data points). 10% Ethanol shows minimal variation in refractive index across the range of temperatures tested while maintaining transparency and viscosity below 0°C (N=3). e) x-z profile of point spread functions measured with water and 10% ethanol immersion medium. Spherical aberrations induced by high refractive index (middle) are equivalent to those seen with a 15 um change in coverglass thickness and can be corrected using the objective collar. Representative data from n=40 repeats.

To prevent condensation at the back of the objective, dry nitrogen was injected into the turret using a 3D-printed insert and a gas line (Figure 2a). To prevent the extra weight of the collar from straining the objective, we designed a 3D-printed support structure to stabilise the objective (Figure 2a). Further details can be found in Figure S4.

A remaining concern for us was the potential heating effect of the sample through energy transfer from the illumination light. We measured the temperature rise in water adjacent to the focal spot of the laser, carefully avoiding direct heating of the temperature probe with the excitation light. A 488 nm laser at 0.448 ± 0.014 mW power caused a maximum of 0.067°C temperature change (Figure 2c). The difference in temperature when the laser was on or off was too small to warrant further efforts to mitigate effects of laser induced heating of the sample.

Finally, the correct choice of immersion medium is critical to obtain optimal resolution and light collection efficiency. Traditional immersion media, such as silicon- and mineral-oils, become viscous when cooled down, hindering movement of the sample over the objective. Below 0°C water is unsuitable for imaging because of freezing (Figure S5). Adding salts to the water would only lower the freezing point by a couple of degrees and is likely to erode the protective coating of the objective. A suitable immersion medium for this system needs to combine the qualities of having a freezing point below 0°C and a refractive index matching that of water over the range of temperatures the system is operated over. Furthermore, the liquid needs to be chosen to avoid damaging the coatings of the objective. Commercially available solutions of water-matching oil [14] were compared to water, culture media, and water based dilutions of ethanol. A 10% dilution of ethanol was found to be the best candidate, yielding a refractive index of 1.34 at 565nm and 0°C (Figure 2d), with a freezing point of -4°C. The refractive index of this mixture was found to stay nearly constant over the visible wavelength range (Figures S6a, b and c). The refractive index of 1.34 is 0.0045 away from the optimum for the lens (namely n of water). This does change the PSF shape and introduces aberrations, but these can be fully corrected for by changing the correction collar to +0.015 mm of glass, making 10% ethanol a suitable immersion medium (Figure 2e).

## 4. Performance of the system

We next assessed whether microscope resolution was affected by mechanical distortion or movement of components caused by our modifications. We compared the performance of the original system without modifications, and that of the instrument with successive additions of the collar, the nitrogen flow, and circulation of the water in the collar. The resolution was assessed by measuring the Full Width at Half Maximum (FWHM) of cross sections of 0.1µm TetraSpeck bead images obtained at 488 nm excitation wavelength using a 60x, NA=1.2 water objective. (Figure 3a). A Student’s t-test (one-tailed, unpaired) was performed, and the results are shown in Figure 3b. No statistically significant differences in resolution were seen to occur (p<0.25 for all cases).

**Figure 3.**
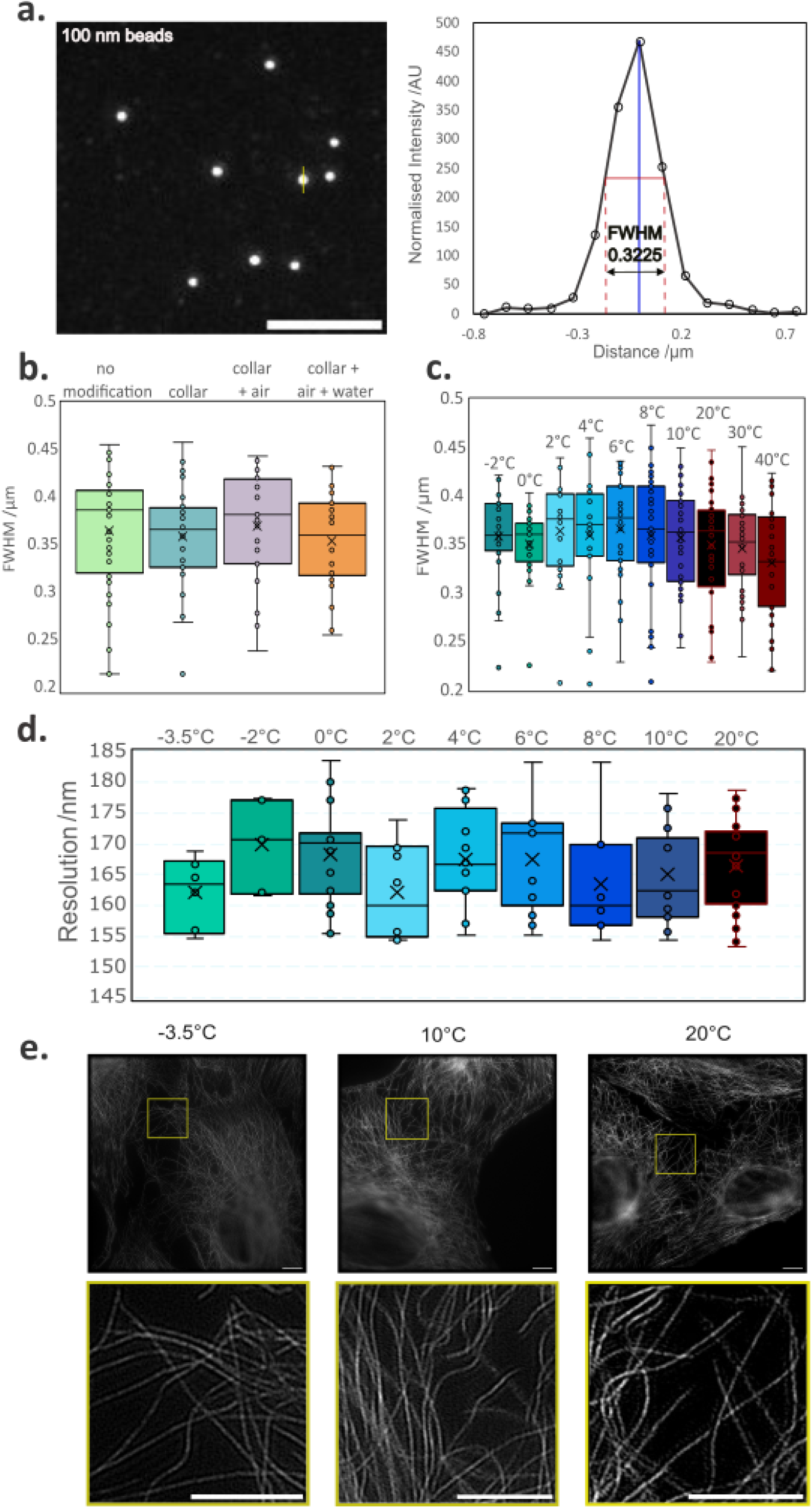
The system performs with near diffraction limited resolution across a 20-degree temperature range. a) Diffraction-limited image of bead monolayer (left) and full-width at half maximum plot of indicated bead (right). No significant differences in PSF size were observed under different system configurations (one-tailed unpaired Student’s t-test, N=30). c) PSF size remains consistent across a 40°C to -2°C temperature range (one-tailed unpaired Student’s t-test, N=26). d) Image resolution estimates from single-image Fourier Ring Correlation (FRC). Estimated resolution stays comparable across the whole temperature range (N=7). e) Reconstruction of SIM images from fixed microtubules at 20°C, 10°C and -3.5°C. Scale bar is 5 µm for all images.

As the temperature of the objective decreases, we expected two concurrent phenomena to affect the resolution. One is a slight worsening of resolution due to the thermal contraction of the objective. Objective barrels are made of brass [15], which has a linear coefficient of thermal expansion (α) of ca. 19×10^−6^K^-1^. Cooling the objective to -20°C could thus generate a contraction of up to 30 µm over the length of the barrel, which is likely to introduce optical aberrations. Lenses within the objective consist of a mix of Flint Glass (α = 8.9×10^−6^ K^-1^) and borosilicate crown glass with a coefficient α = 3.3×10^−6^ K^-1^ [16]. This order of magnitude difference between the material’s thermal expansion coefficient is likely to generate differential contraction as temperatures go lower and is expected to introduce aberrations at very low temperatures.

The other phenomenon is the increase in image contrast afforded at low temperature: the brightness and photostability of the fluorophores used were seen to increase with lowering temperatures, an effect reported in the literature both for fusion proteins and for synthetic fluorophores [17], [18], [19]. Lower temperatures decrease the rotational degrees of freedom available to the molecule, facilitating electron delocalisation and fluorophore brightness. There is furthermore less effect of collisional quenching and other non-radiative processes. [20]

Overall, however, we observed no significant change in optical performance compared to control conditions (Figure 3). The FWHM of obtained point spread functions were recorded at temperatures ranging from 40°C to -2°C using 0.1µm TetraSpec beads at an excitation wavelength of 488 nm and using a 60x water objective. The averages of several PSFs were plotted (Figure 3c), and the corresponding averages for the FWHM were compared for each temperature. Temperatures between -2°C and 40°C were compared. Two temperature points above room temperature (RT) were included to compare the performance of cold microscopy to that of heating the objective and the sample. While heating the objective and sample to 37°C is a common practice in long-term imaging of live mammalian cells [21], [22], [23] the impacts on optical properties of heating the objective 17°C above room temperature is rarely considered. The differences in FWHM were not significant for the PSFs obtained at low temperatures, as p-values for over the entire range tested were above 0.113 when compared to the RT control (one tailed, unpaired Student’s t-test). Cross-comparison of all the conditions highlighted the only significant difference between conditions to be between the 40°C and the cold temperatures (−2°C, 2°C, 6°C and 8°C). Across the cold temperatures, there were no statistical differences (Supplementary Table S2).

To demonstrate the suitability of the method for super-resolution imaging, and to further test the impact on resolution, Fourier Ring Corelation (FRC) [24], [25] was used to assess the resolution of Structured Illumination Microscopy (SIM) data obtained at low temperature. We used fixed samples for this experiment. The resolution was calculated using the FRC method developed by Koho et al. [26] and setting the threshold criteria to ‘half-bit’. Images were taken of microtubules at -3.5°C, -2°C, 0°C, 2°C, 4°C, 6°C, 8°C, 10°C, and at room temperature (Figure 3). The order of recorded temperatures was randomised and changed across repeats to reduce the chances of bleaching impacting the measurements. The Fourier resolution change was insignificant across the temperature range, with p-values for the low temperature data above 0.134 in comparison to the room temperature result (one tailed, unpaired Student’s t-test). One reason for the apparent stability of imaging performance could be that the worsening effect of optical aberrations is offset by improvements in the increasing signal to noise ratio when fluorophores are imaged at lower temperature. The latter appears supported by the greater statistical difference observed in the higher temperature result compared to the low temperature case (Supplementary Table S2).

## 5. Application

Cold-adapted organisms are one example of biological samples for which low-temperature live imaging is a necessity. Antarctic marine fauna has adapted to a very stable and narrow thermal environment. Thermal tolerance studies on such psychrophilic species reveal a 50% failure in essential biological functions at temperatures in the 2-3°C range [2], [27], [28], [29]. To illustrate the need for a cold imaging system, we have chosen to visualise cells from the Antarctic plunder fish *Harpagifer antarcticus*, which is endemic to the Southern Ocean [30]. The entire lifecycle of *H. antarcticus* occurs along the Antarctic Peninsula. Temperatures in this environment typically range only between -1.9°C and 2°C. Although it is possible to acclimate this species, at least short-term, to 3°C [31], this fish, along with other Notothenioids in the Southern Ocean, has an upper lethal thermal threshold of 5-6°C. [32], [33]. We have developed cell culture protocols for *H. antarcticus* but were not previously able to study the cells under physiological conditions due to the lack of suitable optical equipment. Therefore, the cellular mechanisms of adaptation to the cold and consequential molecular and organelle responses to warming in this species, or any polar marine organism, are largely unknown.

To demonstrate that our method improves sample viability during imaging, we compared the impacts of warming in these thermally sensitive samples by imaging at 20°C on a conventional system and at 2°C using the methods presented here. Primary cell cultures (gonad) from *H. antarcticus* were grown at 2°C for 20 days. The cells were then stained using MitoTracker Green and imaged at 2°C, as a control observation of cold-adapted cells in physiological conditions. The cells were then left at 20°C for one hour, approximately the duration of a simple *in vivo* imaging session. Samples were then cooled back to 2°C and imaged again. Mitochondria became more rounded and fragmented as a result, highlighting the damage induced by a room-temperature microscopy experiment on these sensitive samples (Figure 4). We noticed especially that the stress response manifests as a change in the balance between mitochondrial fission and fusion, which is a well-described phenotype of mitochondrial stress and apoptosis [34], [35]. Two metrics best describe this change of phenotype: mitochondrial eccentricity and area. Eccentricity is the measure of how round an object is with the parameter ranging from 0, a perfect sphere (no eccentricity), to 1, characterising a line (fully eccentric). Healthy mitochondria typically display a string-like shape [34]. When the damaged mitochondria become more circular and fragmented, the change in area and eccentricity becomes significant (Figure 4). Student’s t-test between the two conditions show P_area_=0.001 and P_eccentricity_=2.523e^-19^.

**Figure 4.**
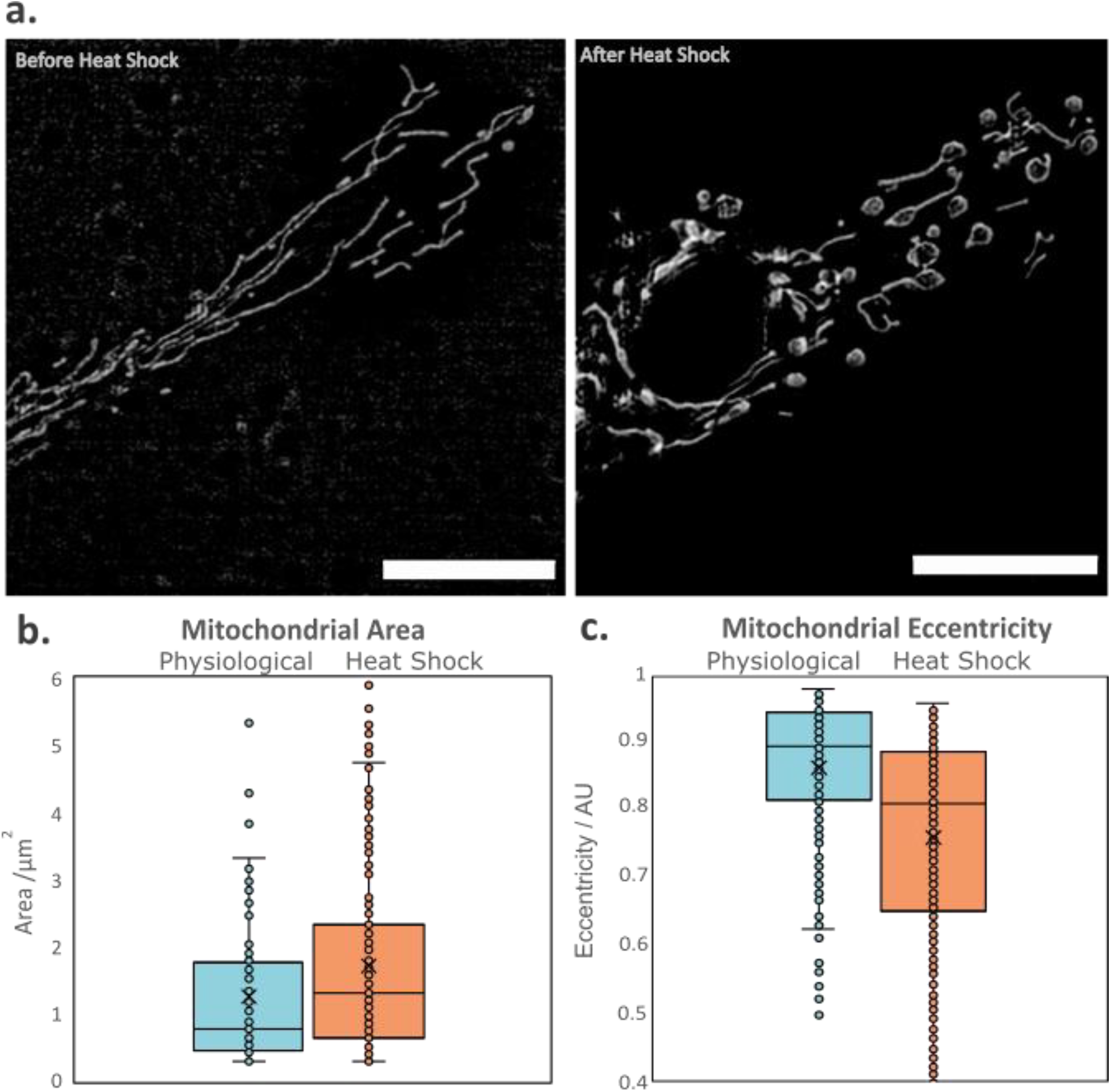
Cold adapted species need to be imaged at their physiological temperature to prevent heat shock. a) Images of mitochondria from cold-adapted cells (primary cell culture of *Harpagifer antarcticus* gonad) acquired at 2°C before (left) and after (right) exposure to room temperature for 1h. Scale bars are 10 µm in both images. b) area and c) eccentricity of the mitochondria in physiological condition pre- and post- temporary exposure to room temperature. n=19, N=3. A significant change in mitochondrial morphology associated with heat shock is observed even after cells have returned to physiological temperatures. This indicates that even short imaging sessions on a non-cooled microscope would perturb natural homeostasis and prevent meaningful *in vivo* study.

The cells were left to recover for 24 hours at 2°C and were imaged the next day, but none survived. This further illustrates how acute heat shock, even when temporary, can irreversibly damage cold-sensitive samples.

## 6. Conclusion

This work proposes a design for a cooling module that enables high-resolution and super-resolution microscopy of samples maintained at, or near, 0°C, and permits observations to be made in live cells. The hardware modifications required to adapt a traditional imaging system for this purpose are relatively minor and our design is compatible with a range of microscope system. The method can thus readily be adopted by other researchers interested in studying biological systems adapted to the cold. We have demonstrated the application of our method for live and super-resolution imaging live cells derived from the Antarctic fish *H. antarcticus*. This opens possibilities to observe in real time mechanisms of proteostasis such as protein folding, denaturation, or aggregation. Further applications are envisaged also for biophysical inquiries into liquid-liquid phase transitions at different temperatures and in various areas of biotechnology and in the food industry. Examples include cold storage stability of cells, tissues or reagents, and transport of transplanted organs, or countless other questions where a microscopic understanding of the effects of cooling and temperature cycling is of interest.

## Supporting information

Supplemental material FigS&-S6 and TablesS1-2

## 7. Acknowledgements

We would like to acknowledge Alec Laws from Instec for his work on the cooling collar.

We would like to thank Elliot Reed for his help in designing and manufacturing of the pieces, Luca Mascheroni for the fixed Vero samples, Meng Lu for his help with the organelle staining, the Rothera marine team for provision of *Harpagifer antarcticus* and The BAS Aquarium Manager for animal husbandry.

This work was supported by the Vice Chancellors Award and School of Technology, University of Cambridge, and the Engineering and Physical Sciences Research Council Centre for Doctoral Training in Sensor Technologies for a Healthy and Sustainable Future [EP/S023046/1] (APMM) ; the EPSRC Centre for Doctoral Training in Connected Electronic and Photonic Systems (JRL), Infinitus Ltd, China (FWvT); Engineering and Physical Sciences Research Council (EP/H018301/1, EP/L015889/1) (ENW) ; Medical Research Council (MR/K015850/1, MR/K02292X/1); Wellcome Trust (089703/Z/09/Z, 3-3249/Z/16/Z) (CFK); and UKRI-NERC core funding to the British Antarctic survey (MSC and LSP).

## 8. Ethics Statement

The Fish used for this study were collected under permit (BAS-S7-2022/01) granted under Section 7 of the Antarctic Act 1994

For the purpose of open access, the authors have applied a Creative Commons Attribution (CC BY) licence to any Author Accepted Manuscript version arising from this submission.

## 9. Author statement

**Anne-Pia M Marty**: Methodology, Investigation, Formal analysis, Visualization, Writing - Original Draft, **Edward N Ward**: Conceptualization, Software, Writing – Review and Editing, **Jacob R Lamb**: Software, **Francesca W van Tartwijk**: Writing - Review & Editing, **Lloyd S Peck**: Supervision, Writing – Review and Editing, **Melody S Clark**: Project conception, Supervision, Funding acquisition, Project administration, Writing - Review & Editing, **Clemens F Kaminski**: Project conception, Supervision, Funding acquisition, Project administration, Writing - Review & Editing. All authors read and approved the final version.

## 1. Tables of reagents, software and equipment

### 1.1 Reagents

| Chemical | supplier | Catalogue number |
| --- | --- | --- |
| MitoTracker green | Invitrogen | M7512 |
| 16% Paraformaldehyde | Thermo Scientific | 28906 |
| Triton | Thermo Scientific | 28314 |
| Glutaraldehyde | Sigma Aldrich | G7776 |
| Mouse AB13120 β-tubulin | Abcam | AB13120 |
| Rabbit AB6046 Antibody | Abcam | AB6046 |
| 0.1μm TetraSpec | Invitrogen | T7279 |

### 1.2 Software

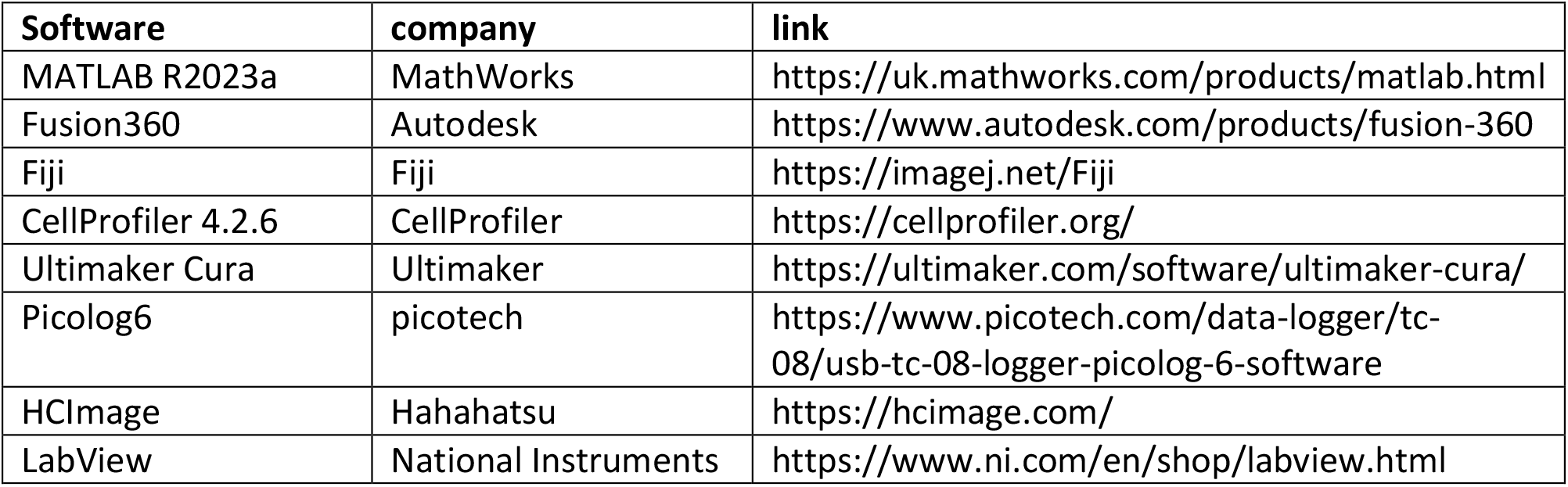

### 1.3 Equipment

| equipment | company | Catalogue number |
| --- | --- | --- |
| thermocouple | Pico Technologies | SE001 |
| Abbe '60' refractometer | Abbe technologies | / |
| LED light source | Thorlabs | LED4D245 |
| LED driver | Thorlabs | DC4104 |
| 3D printer | Ultimaker | UMS3 |
| CNC mill | XYZ Machine tools | 500LR |
| Power meter | Thorlabs | PM100D |
| Power probe | Thorlabs | S130C |
| Microscope stage | Olympus | IX71 |
| 488 laser | Toptica | iBEAM-SMART-488 |
| 561 laser | Coherent | OBIS 561 |
| 640 laser | Cobolt | MLD 640 |
| Spatial light modulator | Forth Dimension Displays | SXGA-3DM |
| Pockels cell | Cornoptics | M350-80-01 |
| Camera | Hamamatsu | C11440 |

## References

[1] A. Clarke, « Ten principles of thermal ecology», in Principles of Thermal Ecology: Temperature, Energy, and Life, A. Clarke, Éd., Oxford University Press, 2017, p. 0. doi: 10.1093/oso/9780199551668.003.0017.

[2] S. J. Hawkins, A. J. Evans, A. C. Dale, L. B. Firth, et I. P. Smith, Éd., « Oceanography and Marine Biology: An Annual Review», in Oceanography and Marine Biology, Taylor & Francis, 2018. doi: 10.1201/9780429454455.

[3] L. S. Peck, P. Convey, et D. K. A. Barnes, « Environmental constraints on life histories in Antarctic ecosystems: tempos, timings and predictability », Biol. Rev., vol. 81, no 1, p. 75–109, févr. 2006, doi: 10.1017/S1464793105006871.

[4] « Linkam stages for hot stage microscopy », Linkam Scientific. Consulté le: 2 mai 2024. [En ligne]. Disponible sur: https://www.linkam.co.uk/general-heating-cooling

[5] « HEATING & COOLING STAGES Precision Thermal stages for Microscopes and Optical Equipment Instec Inc ». Consulté le: 2 mai 2024. [En ligne]. Disponible sur: https://instec.com/portal/list/index/id/49.html

[6] « Heating and Cooling Stages », Analytical technology - Microptik. Consulté le: 2 mai 2024. [En ligne]. Disponible sur: https://www.microptik.eu/product/heating-and-cooling-stages

[7] S. Seki et P. Mazur, « Kinetics and activation energy of recrystallization of intracellular ice in mouse oocytes subjected to interrupted rapid cooling », Cryobiology, vol. 56, no 3, p. 171–180, juin 2008, doi: 10.1016/j.cryobiol.2008.02.001.

[8] A. Guha et R. Devireddy, « Polyvinylpyrrolidone (PVP) Mitigates the Damaging Effects of Intracellular Ice Formation in Adult Stem Cells », Ann. Biomed. Eng., vol. 38, no 5, p. 1826–1835, mai 2010, doi: 10.1007/s10439-010-9963-z.

[9] O. Varisli, H. Scott, C. Agca, et Y. Agca, « The effects of cooling rates and type of freezing extenders on cryosurvival of rat sperm », Cryobiology, vol. 67, no 2, p. 109–116, oct. 2013, doi: 10.1016/j.cryobiol.2013.05.009.

[10] « A Guide to Structured Illumination TIRF Microscopy at High Speed with Multiple Colors ». Consulté le: 10 septembre 2024. [En ligne]. Disponible sur: https://app.jove.com/t/53988/a-guide-to-structured-illumination-tirf-microscopy-at-high-speed-with

[11] R. Heintzmann et T. Huser, « Super-Resolution Structured Illumination Microscopy », Chem. Rev., vol. 117, no 23, p. 13890–13908, déc. 2017, doi: 10.1021/acs.chemrev.7b00218.

[12] M. Müller, V. Mönkemöller, S. Hennig, W. Hübner, et T. Huser, « Open-source image reconstruction of super-resolution structured illumination microscopy data in ImageJ », Nat. Commun., vol. 7, no 1, p. 10980, mars 2016, doi: 10.1038/ncomms10980.

[13] M. R. Lamprecht, D. M. Sabatini, et A. E. Carpenter, « CellProfilerTM: free, versatile software for automated biological image analysis », BioTechniques, vol. 42, no 1, p. 71–75, janv. 2007, doi: 10.2144/000112257.

[14] C. Labs, « Matching Liquids – Cargille Labs ». Consulté le: 23 mai 2024. [En ligne]. Disponible sur: https://www.cargille.com/matching-liquids/

[15] R. Chandler, « The Anatomy of an Objective Lens ». Consulté le: 8 mai 2024. [En ligne]. Disponible sur: https://www.olympus-lifescience.com/en/discovery/the-anatomy-of-an-objective-lens/

[16] H. Golnabi, « COMPARISON OF DIFFERENT GLASS COMPOUNDS FOR INTRINSIC FIBER OPTIC TEMPERATURE SENSORS ».

[17] E. J. Bowen et J. Sahu, « The Effect of Temperature on Fluorescence of Solutions », J. Phys. Chem., vol. 63, no 1, p. 4–7, janv. 1959, doi: 10.1021/j150571a003.

[18] T. M. H. Creemers, A. J. Lock, V. Subramaniam, T. M. Jovin, et S. Völker, « Three photoconvertible forms of green fluorescent protein identified by spectral hole-burning », Nat. Struct. Biol., vol. 6, no 6, p. 557–560, juin 1999, doi: 10.1038/9335.

[19] C. R. Lim, Y. Kimata, M. Oka, K. Nomaguchi, et K. Kohno, « Thermosensitivity of Green Fluorescent Protein Fluorescence Utilized to Reveal Novel Nuclear-Like Compartments in a Mutant Nucleoporin NSP11 », J. Biochem. (Tokyo), vol. 118, no 1, p. 13–17, juill. 1995, doi: 10.1093/oxfordjournals.jbchem.a124868.

[20] P. P. Knox, V. V. Gorokhov, B. N. Korvatovsky, N. P. Grishanova, S. N. Goryachev, et V. Z. Paschenko, « Specific features of the temperature dependence of tryptophan fluorescence lifetime in the temperature range of −170–20 °C », J. Photochem. Photobiol. Chem., vol. 393, p. 112435, avr. 2020, doi: 10.1016/j.jphotochem.2020.112435.

[21] « Okolab - Objective Heater ». Consulté le: 6 mai 2024. [En ligne]. Disponible sur: https://www.oko-lab.com/objective-heater

[22] N. P. Boyer, J.-P. Julien, P. Jung, et A. Brown, « Neurofilament Transport Is Bidirectional In Vivo », eNeuro, vol. 9, no 4, juill. 2022, doi: 10.1523/ENEURO.0138-22.2022.

[23] A. Liu et al., « pHmScarlet is a pH-sensitive red fluorescent protein to monitor exocytosis docking and fusion steps », Nat. Commun., vol. 12, no 1, p. 1413, mars 2021, doi: 10.1038/s41467-021-21666-7.

[24] R. P. J. Nieuwenhuizen et al., « Measuring image resolution in optical nanoscopy », Nat. Methods, vol. 10, no 6, p. 557–562, juin 2013, doi: 10.1038/nmeth.2448.

[25] N. Banterle, K. H. Bui, E. A. Lemke, et M. Beck, « Fourier ring correlation as a resolution criterion for super-resolution microscopy », J. Struct. Biol., vol. 183, no 3, p. 363–367, sept. 2013, doi: 10.1016/j.jsb.2013.05.004.

[26] S. Koho, G. Tortarolo, M. Castello, T. Deguchi, A. Diaspro, et G. Vicidomini, « Fourier ring correlation simplifies image restoration in fluorescence microscopy », Nat. Commun., vol. 10, no 1, p. 3103, juill. 2019, doi: 10.1038/s41467-019-11024-z.

[27] L. S. Peck, K. E. Webb, et D. M. Bailey, « Extreme sensitivity of biological function to temperature in Antarctic marine species », Funct. Ecol., vol. 18, no 5, p. 625–630, 2004, doi: 10.1111/j.0269-8463.2004.00903.x.

[28] L. S. Peck, S. A. Morley, H.-O. Pörtner, et M. S. Clark, « Thermal limits of burrowing capacity are linked to oxygen availability and size in the Antarctic clam Laternula elliptica », Oecologia, vol. 154, no 3, p. 479–484, déc. 2007, doi: 10.1007/s00442-007-0858-0.

[29] H. O. Pörtner, L. Peck, et G. Somero, « Thermal limits and adaptation in marine Antarctic ectotherms: an integrative view », Philos. Trans. R. Soc. B Biol. Sci., vol. 362, no 1488, p. 2233–2258, mai 2007, doi: 10.1098/rstb.2006.1947.

[30] M. G. White et P. J. Burren, « Reproduction and larval growth of Harpagifer antarcticus Nybelin (Pisces, Notothenioidei) », Antarct. Sci., vol. 4, no 4, p. 421–430, déc. 1992, doi: 10.1017/S0954102092000622.

[31] J. M. Navarro, K. Paschke, A. Ortiz, L. Vargas-Chacoff, L. M. Pardo, et N. Valdivia, « The Antarctic fish Harpagifer antarcticus under current temperatures and salinities and future scenarios of climate change », Prog. Oceanogr., vol. 174, p. 37–43, mai 2019, doi: 10.1016/j.pocean.2018.09.001.

[32] « Temperature Tolerance of Some Antarctic Fishes ». Consulté le: 6 mai 2024. [En ligne]. Disponible sur: https://www.science.org/doi/10.1126/science.156.3772.257

[33] K. P. P. Fraser, L. S. Peck, M. S. Clark, A. Clarke, et S. L. Hill, « Life in the freezer: protein metabolism in Antarctic fish », R. Soc. Open Sci., vol. 9, no 3, p. 211272, mars 2022, doi: 10.1098/rsos.211272.

[34] J. R. Friedman et J. Nunnari, « Mitochondrial form and function », Nature, vol. 505, no 7483, p. 335–343, janv. 2014, doi: 10.1038/nature12985.

[35] R. J. Youle et M. Karbowski, « Mitochondrial fission in apoptosis », Nat. Rev. Mol. Cell Biol., vol. 6, no 8, p. 657–663, août 2005, doi: 10.1038/nrm1697.

