## Supplemental material FigS&-S6 and TablesS1-2 for "A high-resolution microscopy system for biological studies of cold-adapted species under physiological conditions"

Figure S1: parameters for thermal simulations

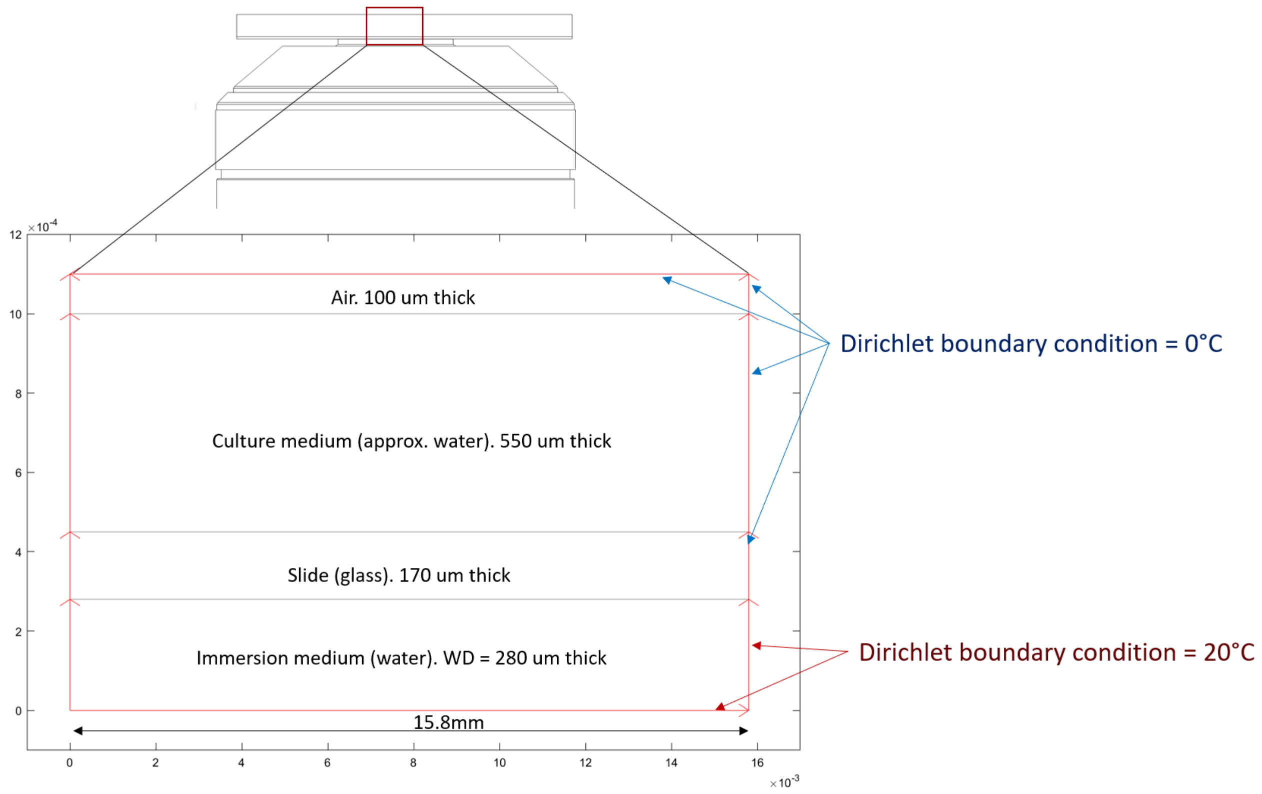


Supplementary Table S1: Parameters used for water, air and glass to compute the thermal models.

| | Culture medium | Immersion Water | Air | Glass |
|---|---|---|---|---|
| Density $\rho$ (kg.m$^{-3}$) | 999.89 | 998.19 | 1.293 | 2530 |
| Heat capacity C (J.kg$^{-1}$.K$^{-1}$) | 4184 | 4184 | 700 | 840 |
| Coefficient of heat conduction k (W.m$^{-1}$.K$^{-1}$) | 0.556 | 0.598 | 0.024 | 0.960 |
| Heat source Q (J.m$^{-2}$.s$^{-1}$) | 0 | 0 | 0 | 0 |
| Convective heat transfer h (W.m$^{-2}$.K$^{-1}$) | 650 | 650 | 55 | 0 |
| External temperature Text (°C) | 0 | 20 | 0 | 0 |

The model was run using the MATLAB PDE toolbox for 10,000 ms for contact with water and 30,000 ms for contact with air.

Figure S2: Renders of the cold incubator used to keep the out-of-focus sample cold. This is used in combination with the objective cooler.

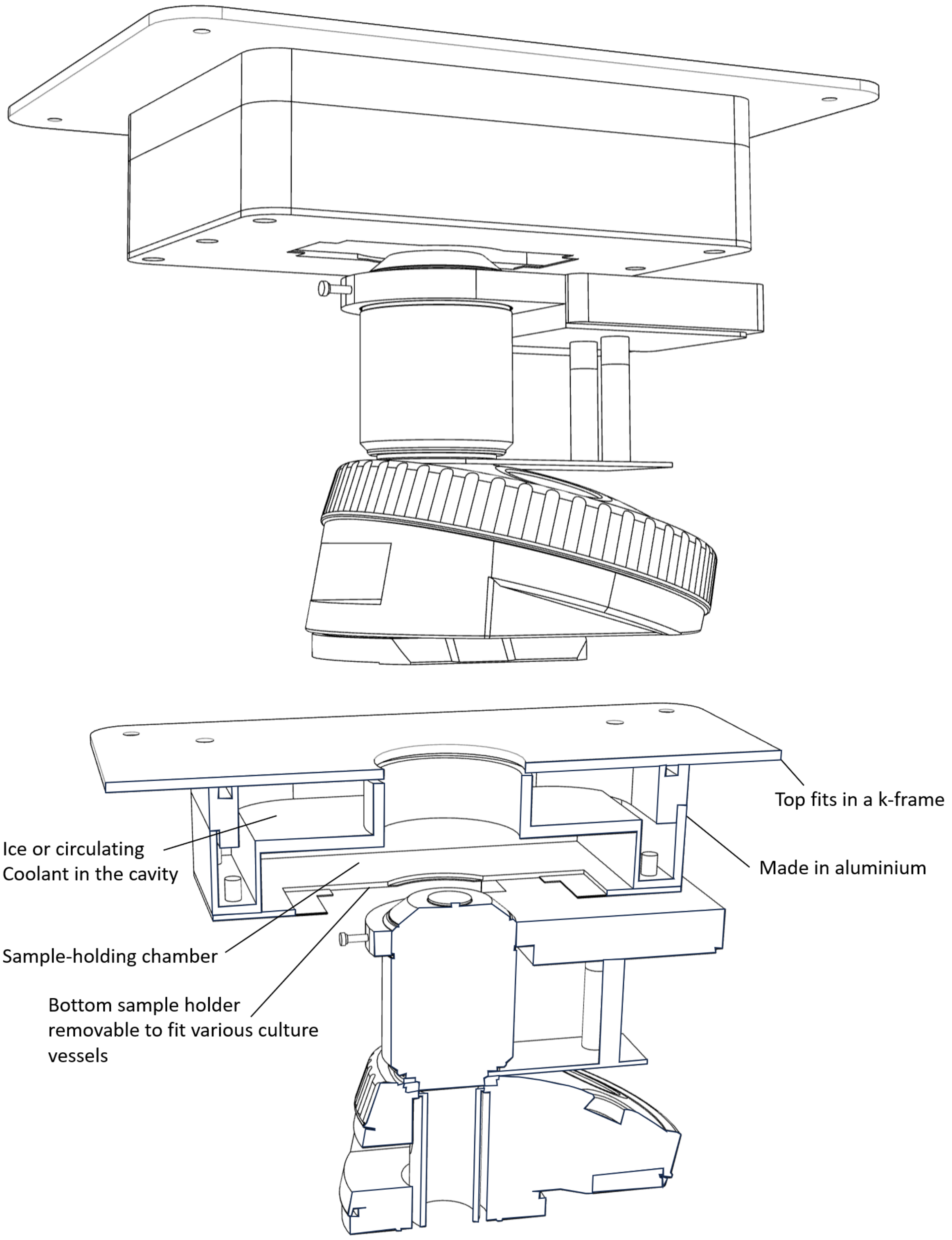

Figure S3: Thermal simulations in Fusion 360 show that a collar cooled to -20 °C brings the temperature in the focal volume down to -6°C.

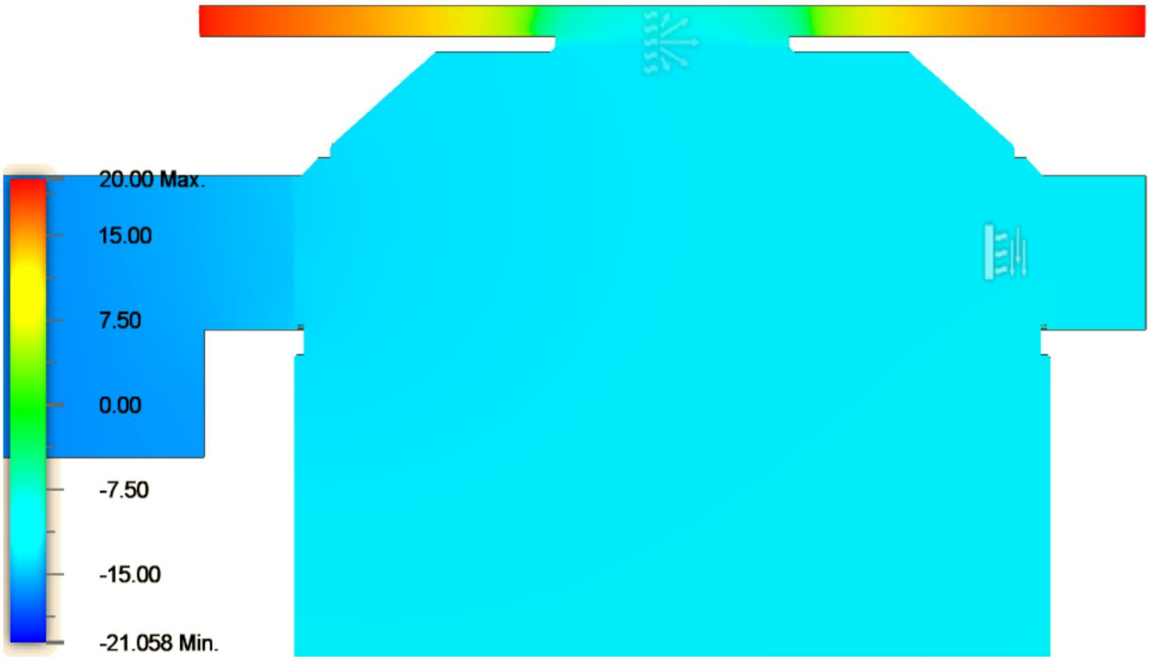

Figure S4: The weight of the collar introduces a noticeable tilt and needs to be offset by a support.

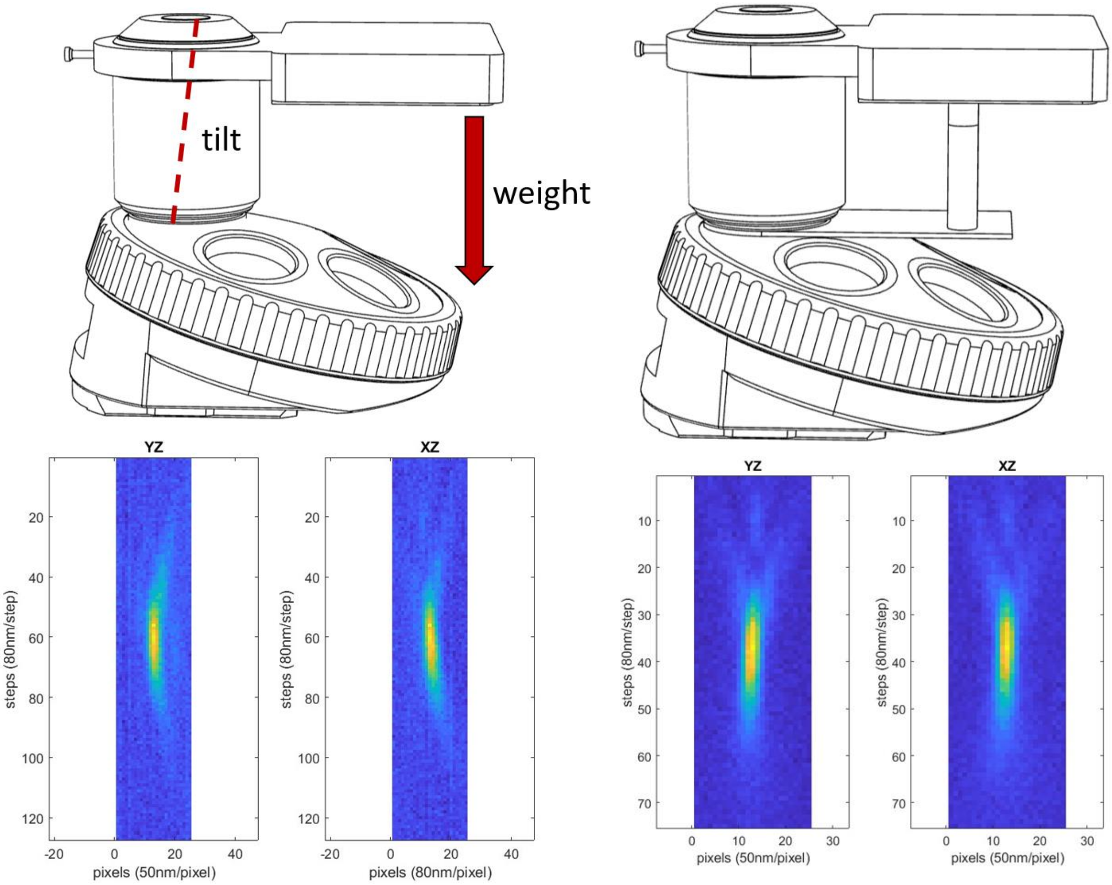

Figure S5 The collar freezes immersion water and a different immersion medium is required.

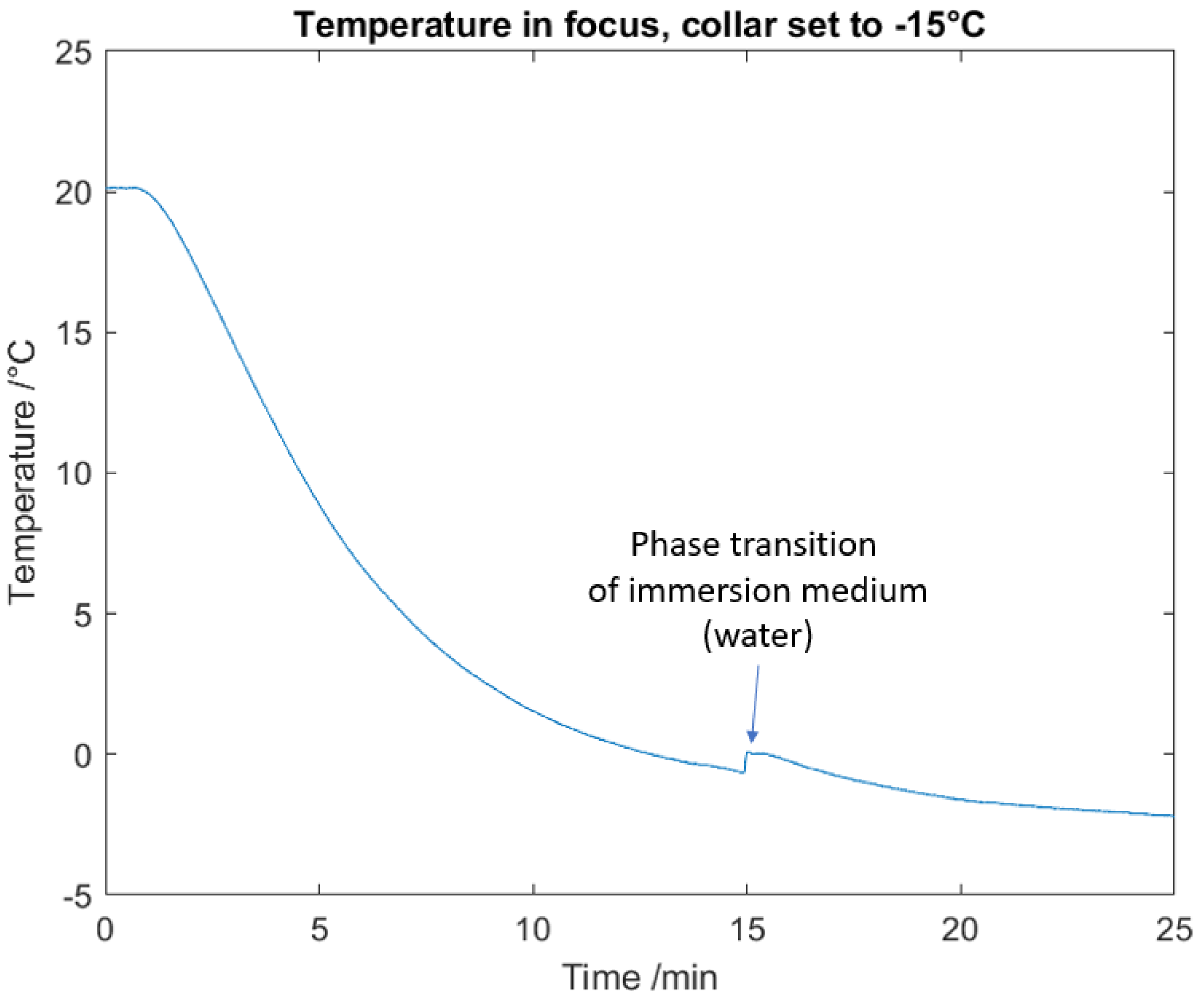

Figure S6: Candidates for sub-0°C immersion media.

a. 565 nm

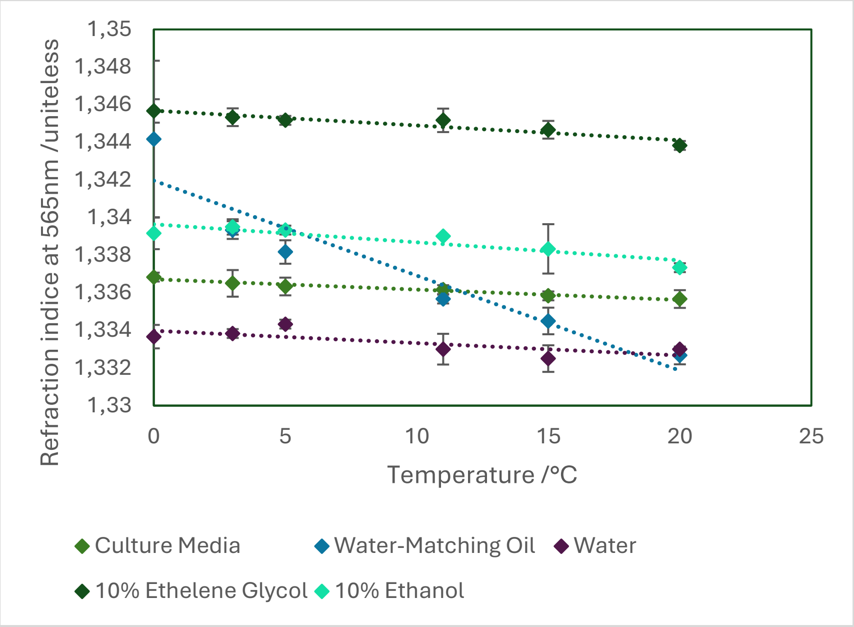


b. 490nm

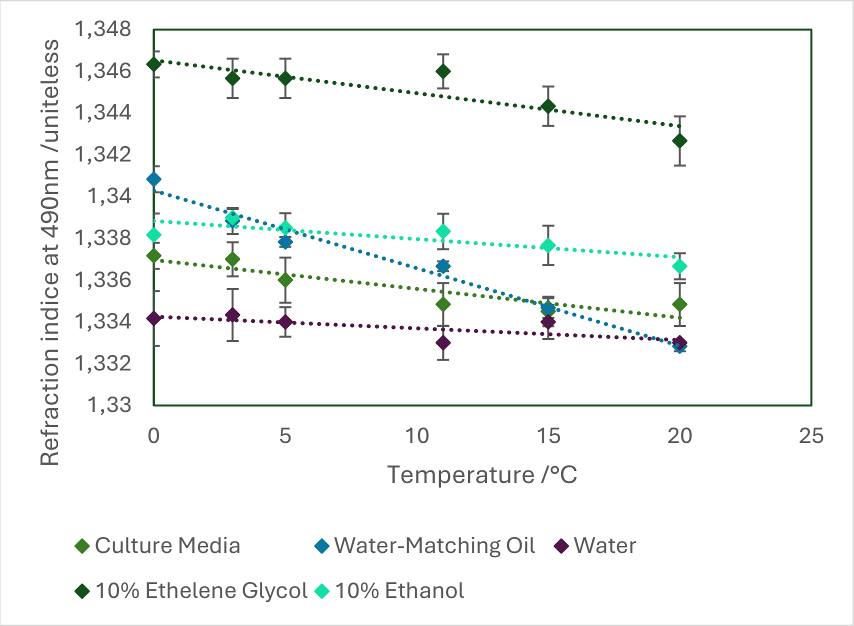

c. 625nm

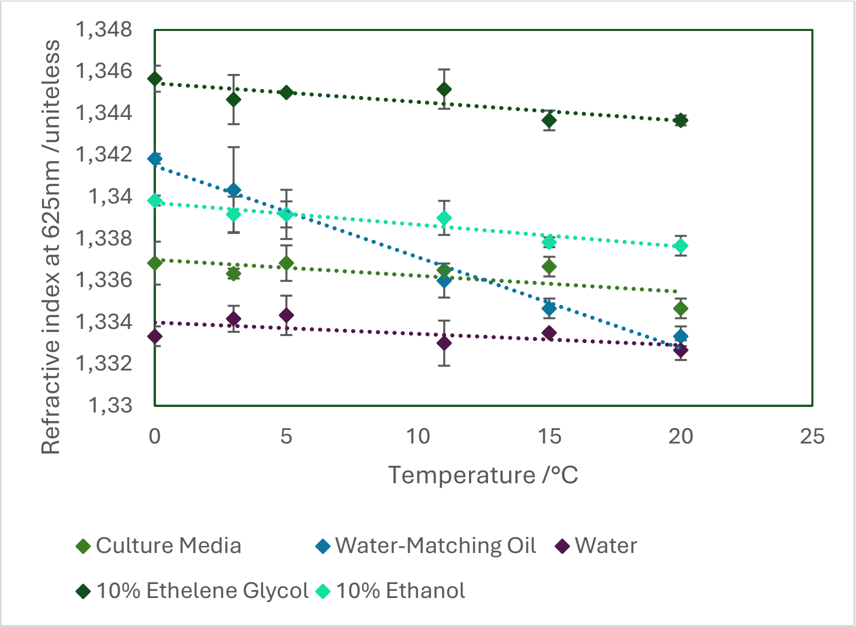

Refractive index at 625nm /uniteless
1,348
1,346
1,344
1,342
1,34
1,338
1,336
1,334
1,332
1,33
0
5
10
15
20
25
Temperature /°C
Culture Media
Water-Matching Oil
Water
10% Ethelene Glycol
10% Ethanol

Supplementary Table 2:

Table of Student's t-tests for the cross-temperature comparisons. The significant differences are highlighted in red.

| | -2°C | 0°C | 2°C | 4°C | 6°C | 8°C | 10°C | 20°C | 30°C | 40°C |
|---|---|---|---|---|---|---|---|---|---|---|
| -2°C | 1 | 0,249301 | 0,337341 | 0,471699 | 0,291509 | 0,465564 | 0,476021 | 0,253385 | 0,163356 | 0,030268 |
| 0°C | | 1 | 0,145255 | 0,264706 | 0,11807 | 0,267216 | 0,278988 | 0,469662 | 0,369785 | 0,092893 |
| 2°C | | | 1 | 0,38549 | 0,452994 | 0,391803 | 0,318685 | 0,156504 | 0,089672 | 0,016638 |
| 4°C | | | | 1 | 0,343437 | 0,494311 | 0,451069 | 0,259323 | 0,356349 | 0,381667 |
| 6°C | | | | | 1 | 0,348916 | 0,273314 | 0,126802 | 0,069113 | 0,011441 |
| 8°C | | | | | | 1 | 0,444117 | 0,249934 | 0,179657 | 0,037725 |
| 10°C | | | | | | | 1 | 0,269311 | 0,182627 | 0,102362 |
| 20°C | | | | | | | | 1 | 0,415602 | 0,112961 |
| 30°C | | | | | | | | | 1 | 0,135449 |
| 40°C | | | | | | | | | | 1 |
